# Synthesis, Biological Activity and Molecular Docking of New Tricyclic Series as α-glucosidase Inhibitors

**DOI:** 10.1101/329557

**Authors:** Hatem A. Abuelizz, Nor Azman N. I Iwana, Rohaya Ahmad, Anouar El-Hassane, Mohamed Marzouk, Rashad Al-Salahi

## Abstract

Diabetes is an emerging metabolic disorder. α-Glucosidase inhibitors, such as acarbose, delay the hydrolysis of carbohydrates by interfering with the digestive enzymes. This action decreases the glucose absorption and the postprandial glucose level. We have synthesized 25 tricyclic 2-phenoxypyrido[3,2-e][1,2,4]triazolo[1,5-a]pyrimidin-5(4H)-ones hybrids and evaluated their α-glucosidase inhibitory activity. Compounds **6h** and **6d** have shown stronger activity than that of acarbose. Compound **6h** exhibited the highest inhibition with an IC_50_ of 104.07 mM. Molecular modelling studies revealed that compound **6h** inhibits α-glucosidase due to the formation of a stable ligand–α-glucosidase complex and extra hydrogen bond interactions, and directed in the binding site by Trp329.

## Introduction

The number of people with diabetes has increased worldwide. According to the global report in diabetes published by WHO in 2014, the number of the diabetic people has increased from 108 million in 1980 to 422 million. Risk factors for diabetes usually include older age, family history, race or ethnicity, obesity and physical inactivity. The prevalence of diabetes among people over 18 years old has increased from 4.7% in 1980 to 8.5% in 2014^1^. Diabetes mellitus is a metabolic disease characterized by hyperglycaemia accompanied by carbohydrate, fat and protein metabolism disturbance. The above-mentioned characteristics of diabetes result from insufficient insulin production in Type I diabetes or ineffective action of insulin in Type II diabetes. The increase in blood sugar levels can lead to serious damage to nerves and blood vessels, which causes a major risk on the individual’s health and quality of life.

Several strategies are used for diabetes management and one of them is controlling postprandial hyperglycaemia. Among postprandial hypoglycaemic agents are α-glucosidase inhibitors. α-Glucosidase inhibitors delay the carbohydrate hydrolysis by inhibiting digestive enzymes, and thus reducing postprandial glucose levels. Moreover, α-glucosidase inhibitors have been found to have a range of biological activities as opposed to other antidiabetic drugs (Acarbose, Miglitol and Voglibose). For example, Celgosivir displays antiviral activity against hepatitis C & B viruses and 1-deoxynojirimycin exhibits anticancer activity (Fig. 1). This variety of biological activity makes α–glucosidase inhibitors a class of compounds with multiple therapeutic applications.

**Figure 1.**
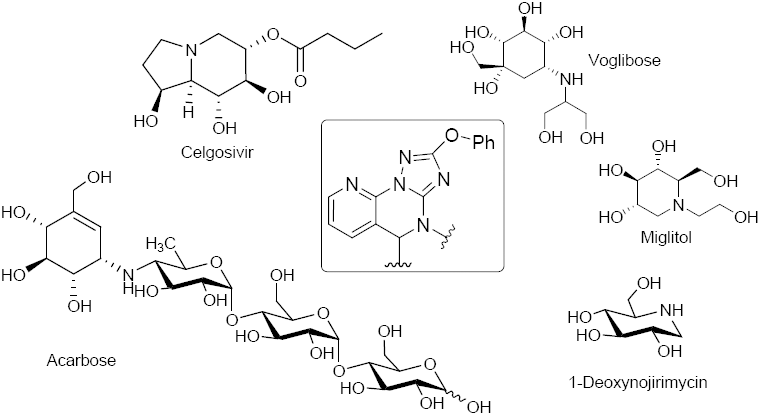
Important clinically used α-glucosidase inhibitors. The boxed structure is the newly synthesized and tested scaffold.

Despite considerable advances in the development of new α-glucosidase inhibitors to treat diabetes, most of the produced compounds were sugar mimetic that requires tedious synthesis. Therefore, a number of compounds mimicking the monosaccharides or oligosaccharides were discovered. Among them, is the *N-* containing carbasugars such as validamine and valienamine, which have proven to be lead compounds for the development of clinically important agents such as acarbose (Fig. 1)^2^. It has been shown that the valienamine moiety in acarbose mimics the oxocarbenium ion-like transition state that binds to and inhibits α-glucosidase^3^. Therefore, acarbose is an effective inhibitor against glucoamylase, sucrase, dextranase and maltase that prevents the digestion of complex carbohydrates, which are subsequently delivered to the colon. However, bacteria in the colon will digest the complex carbohydrates causing gastrointestinal side effects such as flatulence and diarrhoea. Clinical studies have shown that gastrointestinal side effects are the major cause for reduced patient compliance to the diabetic treatment.

The control of postprandial hyperglycaemia by α-glucosidase inhibitors is clinically effective. Several studies have shown that the addition of acarbose to other oral antidiabetic agents is associated with significant improvements in life expectancy and quality of life, and provides excellent money value over the patient lifetime. However, this advantage of the clinically available α-glucosidase inhibitors is losing its value because of the developed gastrointestinal side effects. Therefore, there is a high demand for effective antidiabetic agents.

We are interested in exploring and developing α-glucosidase inhibitors in an attempt to overcome the side effects associated with sugar mimetic agents. According to our previous results, we decided to modify scaffolds well accepted in medicinal chemistry, such as pyridine, triazole, and pyrimidine. These scaffolds have been incorporated in a variety of compounds that displayed interesting pharmacological activities including antimicrobial^4^, antiviral^5^, anticancer^6^, anti-inflammatory^7^, antioxidant^8^ and anticonvulsant^9^ activities. Moreover, pyrido[3,2-*e*][1,2,4]triazolo[1,5,-*a*]pyrimidin-5(4*H*)-ones have been reported as effective fungicidal agents^10^.

Herein, we designed two series of phenoxypyrido-triazolo-pyrimidinones and evaluated their α-glucosidase inhibitory activity. A significant inhibitory activity was observed for the newly synthesized compounds and IC_50_ values lower than that of acarbose were obtained for two compounds. Moreover, molecular docking studies against α-glucosidase were conducted. We could provide insights of a new lead scaffold for the development of new clinically effective antidiabetic drugs.

### Results & Discussion

### 1. Synthesis of fused pyrimidine derivatives

Figure 2 illustrates the synthetic route towards fused pyrimidine derivatives. The starting material (**A**) was prepared according to a reported procedure^11^. The incorporation of a hydrazine moiety to *N*- cyanoimidocarbonates to generate triazoles has been successfully achieved and is well documented^12^ Thus, 1,2,4-triazole intermediate **B** was obtained when an ethanolic solution of diphenoxy-*N*- cyanoimidocarbonate was allowed to react with **A** in a basic medium (triethylamine) at room temperature. Upon the treatment of **B** with conc. HCl, initially at room temperature followed by heating, afforded novel 2-phenoxypyrido[3,2-e][1,2,4]triazolo[1,5-a]pyrimidin-5(4H)-one (**1**) in 78% yield. The chemical structure of compound **1** was established on the basis of NMR, IR and mass spectra. The IR spectrum of **1** was characterized by a band at 1703 cm^−1^ due to a strong stretching vibration of a C=O group.

**Figure 2.**
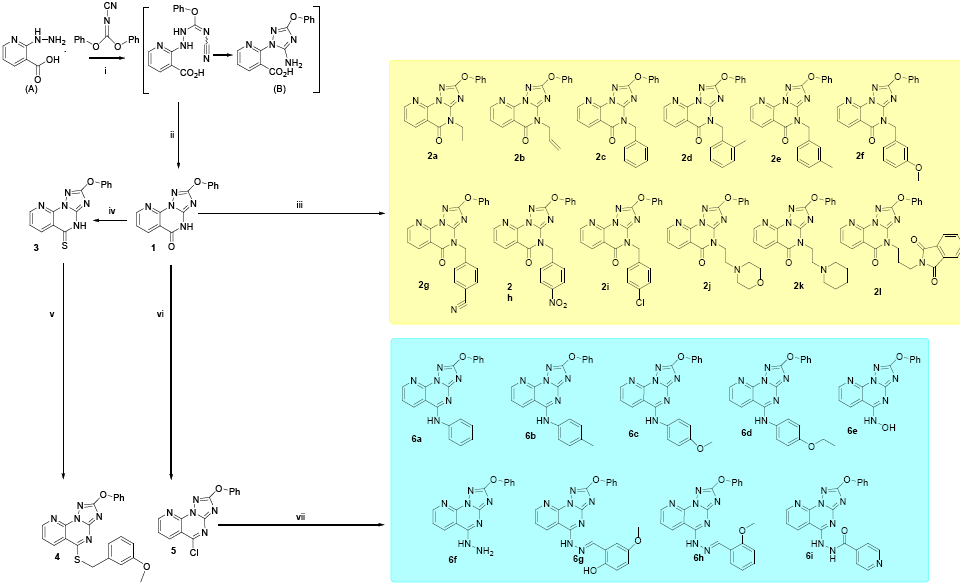
Synthetic routes towards pyrimidine derivatives (**1**–**6**). i: Et_3_N, EtOH, overnight stirring; ii: conc. HCl, 80 °C; iii: DMF, K_2_CO_3_, alkyl or heteroalkyl halides; iv: P_2_S_5_, pyridine; v: DMF, K_2_CO_3_, 3-methoxybenzylbromide 90 °C; vi: POCl_3_, benzene; vii: pyridine, aromatic amines; pyridine, hydroxylamine HCl; toluene, isoniazid; EtOH, hydrazine hydrate; EtOH, glacial acetic acid, aldehydes.

Under suitable conditions^12^, the reaction of compound **1** with alkyl or heteroalkyl halides gave the expected *N*-alkylated pyrido[3,2-e][1,2,4]triazolo[1,5-a]pyrimidin-5(4H)-ones (**2a–l**) in 45-83% yields. The IR and NMR data confirmed the formation of *N*-alkylated products **2a-l** (Figure 2). Compounds **2a-l** were obtained as coloured amorphous powders and their IR spectra displayed absorption bands at 1670-1688 cm^−1^ due to the C=O group. Boiling equimolar amounts of target **1** and phosphorous pentasulfide in absolute pyridine furnished the desired 2-phenoxypyrido[3,2-e][1,2,4]triazolo[1,5-a]pyrimidin-5(4H)-thione (**3**) as a yellow amorphous powder in excellent yield (90%). The IR and ^13^C NMR spectra of compound **3** displays a weak absorption band (1272 cm^−1^) and a peak resonance (186.0 ppm), respectively, due to the C=S group. The conversion of **3** into **4** was carried out in basic medium using benzyl bromide. The transformation of **1** into **5** was successfully achieved employing phosphorous oxychloride as chlorinating agent in boiling benzene followed by treatment with a saturated basic solution. The formation of **5** was followed by NMR and IR spectroscopy; the latter confirmed the completely disappearance of the C=O band characteristic in compound **1** at 1703 cm^−1^. The conversion of the lactam group in **1** into an imidoyl chloride moiety (**5**) was expected to provide a valuable intermediate for further displacement reactions with several nucleophiles. Reaction of **5** with hydroxylamine HCl in a molar ratio of 1:2.5 in boiling pyridine provided the corresponding product **6e** in 63% yield (Figure 2), while treatment of target **5** with several aromatic amines in pyridine in a molar ratio of 1:5 afforded the corresponding compounds **6a-d** in 60-68% yields. The chlorine was replaced in **5** by isoniazid in boiling toluene to give amidrazone **6i** in good yield (74%). Hydrazinolysis of **5** in boiling ethanol produced the corresponding compound **6f**, which upon treatment with equimolar amounts of aldehydes resulted in the formation of hydrazone derivatives **6g** and **6h** in good yields (Figure 2).

The ^1^H-NMR spectra of targets **1**–**6** were characterized by two consistent spin coupling systems, each consisting of three types of protons, indicative of the main 2-phenoxypyrido[3,2-*e*][1,2,4]triazolo[1,5- *a*]pyrimidine structure. The first system consists of a triplet (H-3′/5′, 2H), doublet (H-2′/6′, 2H) and triplet (H-4′, 1H) in the range δ 7.50–7.25 ppm, corresponding to the five protons of the 2-phenoxy moiety. The second one corresponds to the three protons of the pyridine-ring (H-6, H-7 and H-8), mostly in the form of two doublets (H-8 & H-6) and a double of doublets (H-7) in the range of δ 8.9–7.5 ppm (see the experimental data). This main structure of phenoxy-pyrido-triazolopyrimidinone was also clearly interpretable from its 12 characteristic ^13^C-resonances, including five resonances of the fused pyrido ring, four of the phenoxy moiety along with the characteristic three signals of C-2, C-3a and C-5. The successful synthesis of the different groups of products (**1**; **2a–l**; **3**; **4**; **5** or **6a–i**) was separately supported by the characteristic splitting patterns, δ and J values of the newly added moieties (alkyl or aryl) in the corresponding ^1^H and ^13^C NMR spectra. In addition, the change of structural features among all groups was mainly validated in the form of different δ values for the C-5 due to its structural change from C=O (**1** & **2a–l**) to C=S (**3**), C-S-R (**4**), C-Cl (**5**) or C-NHR (**6a–i**) (see the experimental data).

## 2. Biological evaluation

### 2.1. α-Glucosidase inhibitory assay

To explore the potential biological activity of the synthesized pyrimidine fused derivatives (**1**–**6**), *in vitro* screening of α-glucosidase inhibition was performed. The enzyme was treated with equal concentrations (100 µg/mL) of each compound. The test was repeated three times for each compound and the average percent inhibition was obtained. The inhibitory activity of all the tested compounds was between 30% and 63%. The eight compounds (out of the 25 tested) that exhibited more than 50% of the average percent inhibition were further evaluated and their IC_50_ values compared to that of acarbose. Compounds **3**, **6c** and **6d** showed average percent inhibition of more than 60%. On the other hand, compounds **2a**, **2d**, **6a**, **6b** and **6h** showed average percent inhibition of more than 50% but less than 60%.

In this study, we demonstrated that the introduction of a branch to the pyrimidine scaffold through a nitrogen bridge results in a series of synthetic compounds with significant inhibitory activity against α-glucosidase. The open site available for *N*-substitution at the fused pyrido-triazolo-pyrimidine rings is on the nitrogen at position 4. However, these derivatives did not exhibit any inhibitory activity except for compounds **2a** and **2d**. This suggests that the derivatization site has no significant effect on the inhibitory activity. The average percent inhibitions of **2a** and **2d** were 50.36% and 57.48%, respectively. The replacement of the ketone group of the pyrimidinone with amino-bridged substitutes resulted in the production of compounds **6a**–**6i**. Compounds **6a–d** and **6h** exhibited average percent inhibition of more than 50% (Figure 3). The comparison between the average percent inhibition of compounds **6a**–**6i** and **2a**–**2l** suggests that the amino-bridge substitution of the pyrimidine ring at position 4 is responsible for the strong inhibitory activity.

**Figure 3.**
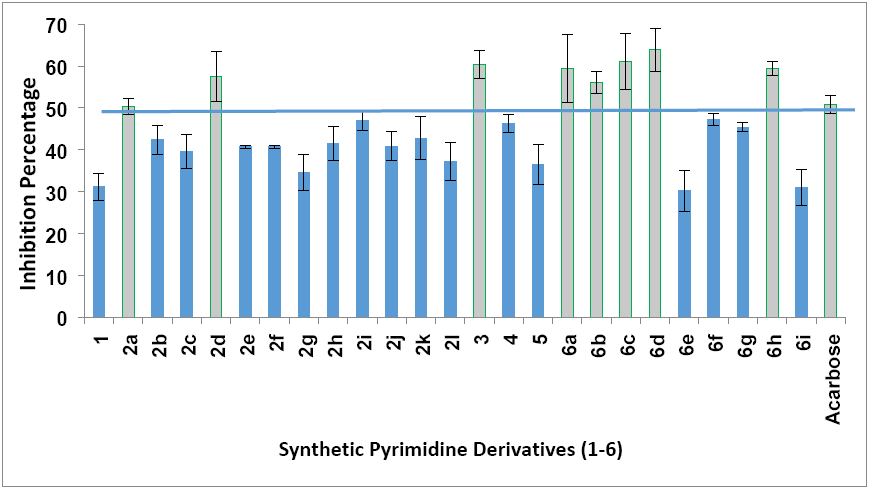
α-Glucosidase inhibitory assay of the synthesized fused pyrimidine derivatives (1–6). The IC_50_ was calculated for compounds that exhibited average percent inhibition of more than 50%.

### 2.2. IC_50_ values of the active compounds

The pharmaceutical parameters of promising inhibitors, including the corresponding IC_50_ values, were calculated. Acarbose, a commonly used antidiabetic drug due to its inhibitory activity against α-glucosidase, was used as positive control. Compound **3** at a concentration of 100 µg/mL exhibited good activity against α-glucosidase enzyme (60.38%). Instead, products derivatized at position 4 has displayed weak inhibitory activity. Only two *N*-alkylated derivatives (ethyl and 2-methylbenzyl) showed more than 50% average percent inhibition (50.36% and 57.48%, respectively). The IC_50_ values of these compounds followed a similar trend; the IC_50_ of compound **3** was 167.82 µM and those of compound **2a** and **2d** were 304.13 µM and 173.83 µM, respectively (Figure 4).

**Figure 4.**
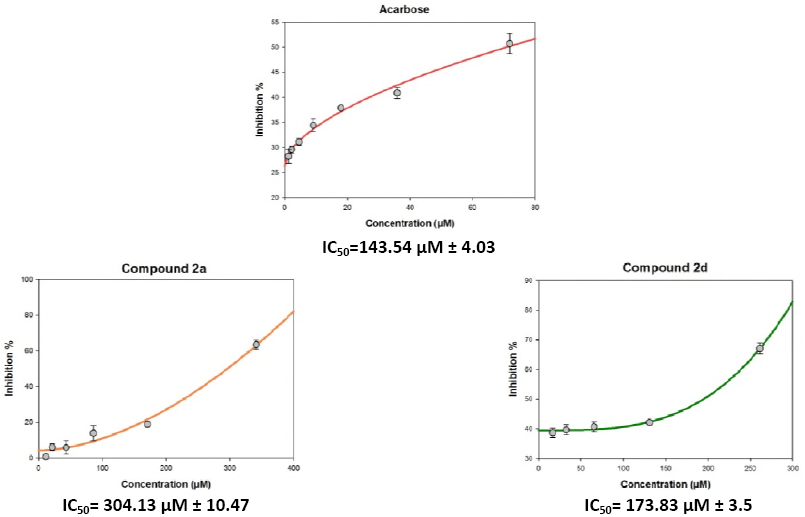
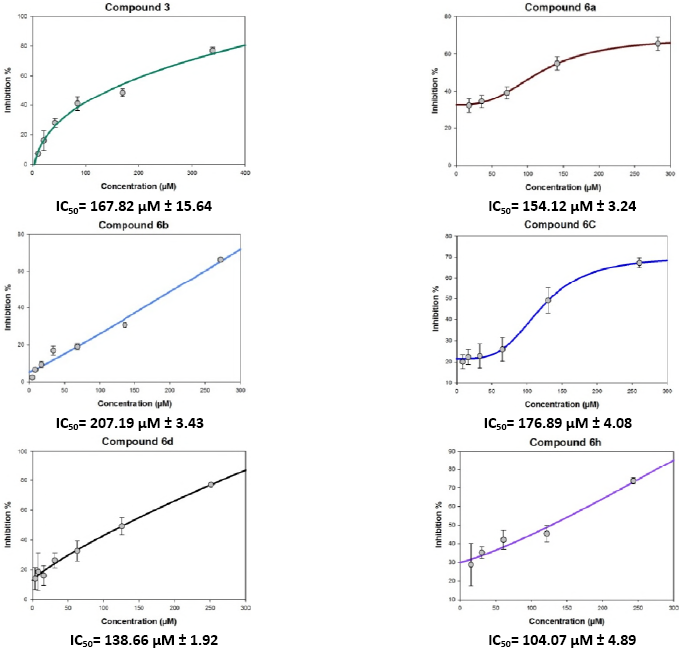
IC_50_ values of **2a**, **2d**, **3**, **6a**, **6b**, **6c, 6d** and **6h** with average percent inhibition of more than 50%. More than five concentrations were evaluated and the inhibitory activity compared to that of acarbose, IC_50_ values ± SEM (P ≤ 0.05).

On the other hand, the replacement of the chloro group in **5** by different nucleophiles provided derivatives with similar or lower IC_50_ values than that of acarbose. This is exemplified by compounds **6a**, **6b**, **6c**, **6d** and **6h**. The rank order of potency according to the IC_50_ values, including acarbose is **6h** > **6d** > acarbose > **6a** > **6c** > **6b**. The 2-methoxy-benzyl-methanimine (IC_50_ **=** 104.07 µM) and 4-ethoxy-aniline (IC_50_ **=** 138.66 µM) substituted derivatives exhibit the best activity when compared to acarbose (IC_50_ =143.54 µM). The IC_50_ values of the amino substituted derivatives ranked as aniline > 4-methoxy-aniline > 4-methyl-aniline. While the IC_50_ value was slightly reduced by the parent aniline compound **6a** (IC_50_ **=** 154.12 µM), the 4-methyl-aniline and 4-methoxy-aniline derivatives showed IC_50_ values of 207.19 and 176.89 µM, respectively.

## 3. Molecular docking

To understand the observed α-glucosidase inhibitory activities of the synthesized compounds, molecular docking calculations were performed to study the binding modes of the target enzyme and docked synthesized derivatives. Therefore, the number of established intermolecular hydrogen bonds, binding energies of stable ligand–α-glucosidase complexes and number of closest amino acid residues surrounding the binding site of the α-glucosidase enzyme were determined for the synthesized compounds (Table 1 and Figure 5).

**Table 1.**
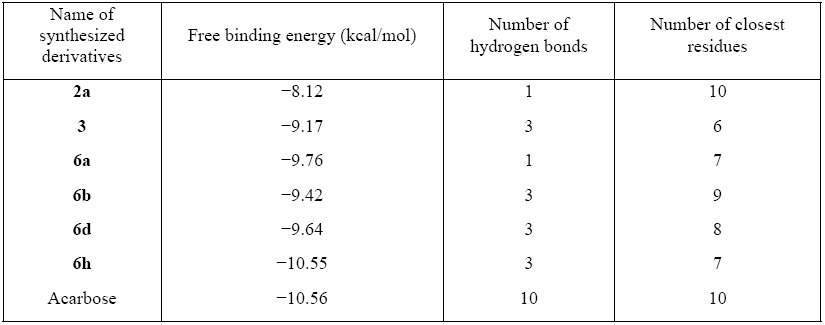
Inhibitory activities and binding energies of docked 2-phenoxypyrido[3,2-e][1,2,4]triazolo[1,5-a]pyrimidin-5(4H)-ones derivatives and acarbose into the active site of α-glucosidase.

| Name of synthesized derivatives | Free binding energy (kcal/mol) | Number of hydrogen bonds | Number of closest residues |
| --- | --- | --- | --- |
| **2a** | -8.12 | 1 | 10 |
| **3** | -9.17 | 3 | 6 |
| **6a** | -9.76 | 1 | 7 |
| **6b** | -9.42 | 3 | 9 |
| **6d** | -9.64 | 3 | 8 |
| **6h** | -10.55 | 3 | 7 |
| Acarbose | -10.56 | 10 | 10 |

**Figure 5.**
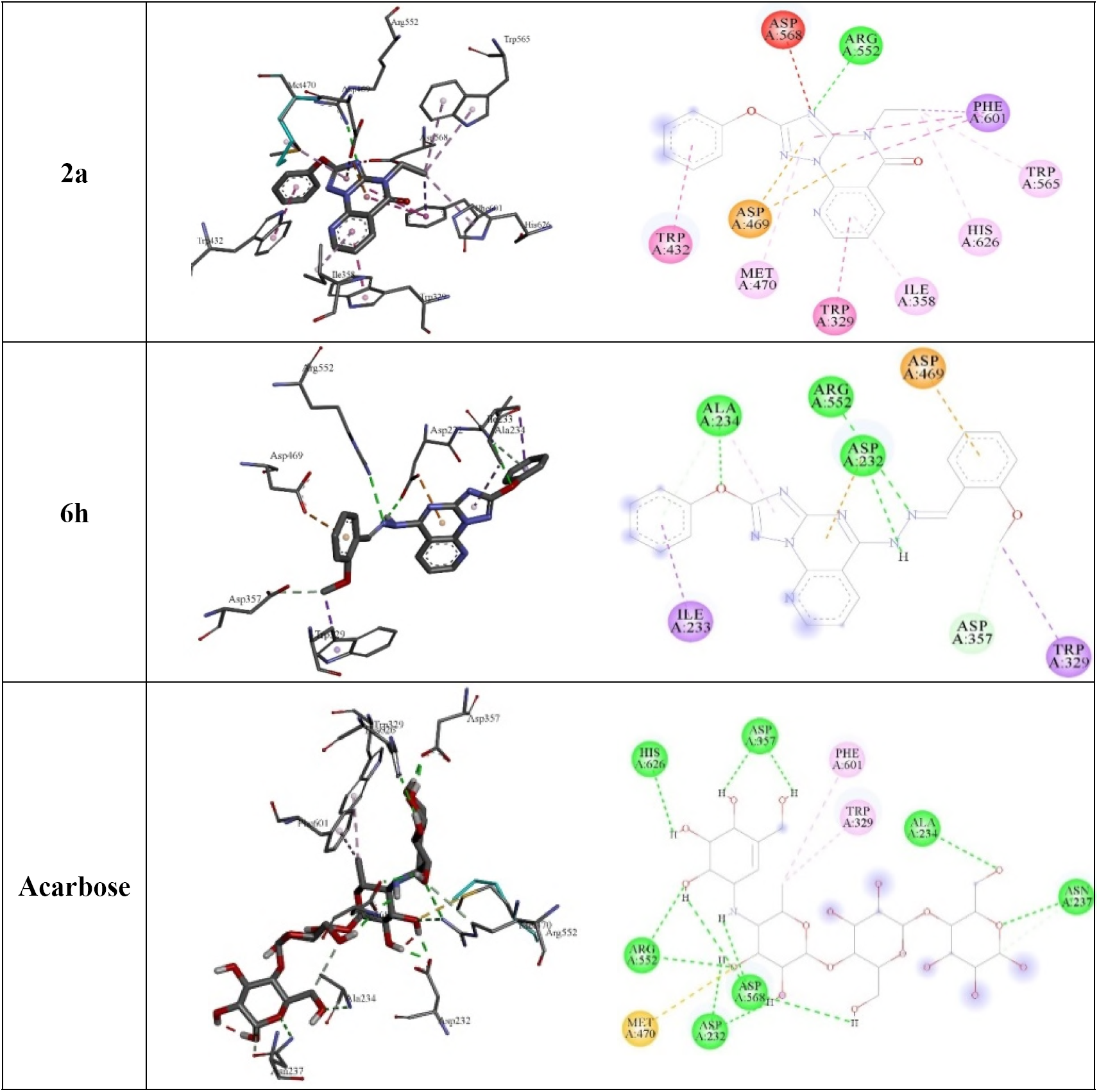
Molecular docking comparison of acarbose with compounds **2a** and **6h**. 3D binding conformation (right) and 2D binding conformation (left) showing the closest interactions between the active site residues of α-glucosidase and the most active (**6h**), least active (**2a**) synthesized derivatives and acarbose.

All the synthesized derivatives formed a complex with the target enzyme. The relative standard deviation of the binding energy of the formed complex was less than 2.34 kcal/mol. From Table 1 and Figure 5, it can be assumed that the maximal inhibition activity of compound **6h** is mainly attributable to its binding energy and the stability of the ligand–α-glucosidase complex. Thus, the binding energy of docked **6h** was found to be the lowest (−10.56 kcal/mol) among the energies of the docked derivatives (Table 1 and Figure 5). Compounds **6a** and **6b** differ from compound **6h** only at the substitution at the amino group (phenyl-, 4-Me-phenyl and *N*-(2-methoxy benzyl)methanimine, respectively; Figure 2). The inhibitory activity of compound **6a** was higher than that of **6b**, and this is may be ascribed to the higher stability of the **6a**–α-glucosidase complex (binding energy = −9.76 kcal/mol) than the **6b**–α-glucosidase complex (binding energy = −9.17 kcal/mol). The number of established hydrogen bonds can be invoked by the reference drug acarbose. The inhibition activity of acarbose correlates with the number hydroxyl groups and the number of the established hydrogen bonds between the ligand and the active site of the α-glucosidase enzyme (Figure 5). Acarbose can establish 10 hydrogen bonds with Asp232, Ala234, Asn237, Asp357, Arg552, Asp568 and His626. Moreover, two hydrophobic interactions are observed between the methyl group of the deoxyglucosamine moiety and residues Phe601 and Trp329. Although compound **2a** can interact with 10 residues at the active site, it can only form one hydrogen bond between the nitrogen group on the triazole ring and Arg552. However, this binding is not stable and provides a free binding energy of −8.12 kcal/mol. Instead, three residues appear to interact with compound **6h**, Ala234, Arg552 and Asp469, to which **6h** can establish hydrogen bonds. These three hydrogen bonds enhance the ligand-α-glucosidase complex stability and reduce the free binding energy of compound **6h** to −10.55 kcal/mol. Trp329 can also form hydrophobic bonds with compounds **2a** and **6h** as in the case of acarbose, which is responsible for the binding conformation of the compounds in the active site.

## Conclusion

We have prepared twenty-five compounds containing fused pyrimidine, pyrido and triazole rings. The synthesized compounds were characterized and subjected to α-glycosidase inhibitory assay. Six promising compounds that showed more than 50% of the average percent inhibition were subjected to further assays to determine their IC_50_. Acarbose was used as reference drug to compare with the synthesized compounds. Compounds **6d** and **6h** showed a significant inhibitory activity against α-glycosidase with lower IC_50_ values than that of acarbose. Molecular docking studies were performed and showed that the number of established intermolecular hydrogen bonds, the binding energies of the stable ligand–α-glucosidase complexes and the number of closest amino acid residues surrounding the binding site are responsible for the inhibitory activity of compound **6h**, which is higher than that of acarbose. Moreover, Trp329 plays a significant role in directing the ligand in the binding site of the α-glucosidase enzyme. This study provides a new structural insight for the development of novel α-glucosidase inhibitors, which differ from the commonly employed carbasugars, and which can eventually be transformed into new clinical drugs for the treatment of type II diabetes.

## Materials and Methods

### Chemistry

Melting points were measured in open-glass capillaries using a STUART SMP 10 melting point apparatus and are uncorrected. NMR spectra were recorder in DMSO-d6 either on a Bruker AMX 500 or 700 spectrometer. Chemical shifts are reported relative to TMS as δ ppm values. Spectrometers operated at either 500 or 700 MHz for ^1^H NMR and 175 MHz for ^13^C NMR. Coupling constants (J values) are given in Hz. High resolution mass spectra (HREI-MS) were measured on a JEOL MStation JMS-700 system. A Perkin Elmer FT-IR Spectrum BX spectrometer was used to record the IR spectra (KBr, v, cm^−1^). TLC on DC Mikrokarten polygram SIL G/UV254 (thickness: 0.25 mm from the Macherey-Nagel Firm Duren) was used to monitor the reactions and check the purity of compounds.

### Procedure for the synthesis of 2-phenoxypyrido[3,2-*e*][1,2,4]triazolo[1,5,-*a*]pyrimidin-5(4*H*)-one (1)

2-Hydrazino-3-carboxylic pyridine (5.5 mmol) was added portion wise to a solution of diphenoxy-*N*-cyanoimidocarbonate (5 mmol) in ethanol (20 mL) at room temperature. Afterwards, triethylamine (20 mmol) was added dropwise over a 10-min period. After the addition was complete, the reaction mixture was left to stir at room temperature overnight. The mixture was acidified with conc. HCl under ice cooling and then heated for 1–2 h at 80 °C. After cooling, the mixture was poured into ice/water, the resulting solid was collected by filtration, washed with water and dried. The product was obtained as a white amorphous powder (78%); mp: 252–254 °C; IR: 1703 (C=O) cm^−1^; ^1^H NMR (700 MHz, DMSO-*d*_*6*_): δ 13.27 (br-s, 1H, NH), 8.82 (dd, J = 4.8, 1.7 Hz, 1H, H-8), 8.55 (dd, J = 7.8, 1.7 Hz, 1H, H-6), 7.59 (dd, J = 7.8, 4.8 Hz, 1H, H-7), 7.47 (t, J = 7.8 Hz, 2H, H-3′/5′), 7.34 (d, J = 7.8 Hz, 2H, H-2′/6′), 7.28 (t, J= 7.8 Hz, 1H, H-4′); ^13^C NMR (175 MHz, DMSO-*d*_*6*_): δ 166.9 (C-2), 160.1 (C-5), 154.8 (C-3a), 154.5 (C-1′), 149.7 (C-9a), 146.7 (C-8), 138.4 (C-6), 130.3 (C-3′/5′), 125.6 (C-4′), 122.4 (C-5a), 120.3 (C-2′/6′), 113.7 (C-7); HRMS m/z (EI): [M]^•+^calcd. for C_14_H_9_N_5_O_2_, 279.0756; found 279.0797.

### General procedure for the synthesis of compounds 2a–l

At room temperature, potassium carbonate (0.6 mmol) was added portion wise over a period of 5 min to a stirred solution of **1** (0.5 mmol) in DMF (5 mL). After 10 min, the appropriate alkyl or heteroalkyl halide (1 mmol) was added dropwise and then the resulting mixture was left stirring at room temperature for 18 h. Afterwards, the mixture was poured into ice/water, the resulting solid was filtered, then washed with water and dried.

### 4-Ethyl-2-phenoxypyrido[3,2-*e*][1,2,4]triazolo[1,5,-*a*]pyrimidin-5(4*H*)-one (2a)

White amorphous powder (73%); mp: 173–175 °C; IR: 1681 (C=O) cm^−1^; ^1^H NMR (500 MHz, DMSO-*d*_*6*_): δ 8.84 (dd, J = 4.8, 1.7 Hz, 1H, H-8), 8.61 (dd, J = 7.8, 1.7 Hz, 1H, H-6), 7.61 (dd, J = 7.8, 4.8 Hz, 1H, H-7), 7.49 (t, J = 7.8 Hz, 2H, H-3′/5′), 7.37 (d, J = 7.8 Hz, 2H, H-2′/6′), 7.30 (t, J = 7.8 Hz, 1H, H-4′), 4.16 (q, J = 7.2 Hz, 2H, H-1′′), 1.31 (t, J = 7.2 Hz, 3H, H-2′′); ^13^C NMR (175 MHz, DMSO-*d*_*6*_): δ 166.6 (C-2), 158.9 (C-5), 154.8 (C-3a), 154.4 (C-1′), 150.4 (C-9a), 145.9 (C-8), 138.7 (C-6), 130.4 (C-3′/5′), 125.7 (C-4′), 122.5 (C-5a), 120.1 (C-2′/6′), 113.1 (C-7), 39.2 (C-1′′), 12.8 (C-2′′); HRMS *m/z* (EI): [M]^•+^ calcd. for C_16_H_13_N_5_O_2_, 307.1069; found 307.1098.

### 4-Allyl-2-phenoxypyrido[3,2-*e*][1,2,4]triazolo[1,5,-*a*]pyrimidin-5(4*H*)-one (2b)

White amorphous powder (78%); mp: 170–172 °C; IR: 1686 (C=O) cm^−1^; ^1^H NMR (700 MHz, DMSO-*d*_*6*_): δ 8.85 (dd, J = 4.8, 1.7 Hz, 1H, H-8), 8.62 (dd, J = 7.8, 1.7 Hz, 1H, H-6), 7.63 (dd, J = 7.8, 4.8 Hz, 1H, H-7), 7.49 (t, J = 7.8 Hz, 2H, H-3′/5′), 7.37 (d, J = 7.8 Hz, 2H, H-2′/6′), 7.31 (t, J = 7.8 Hz, 1H, H-4′), 5.97 (m, 1H, H-2′′), 5.30 (d, J = 17.2 Hz, 1H, H-3a′′), 5.20 (d, J = 10.4 Hz, 1H, H-3b′′), 4.73 (d, J = 4.9 Hz, 1H, H-1′′); ^13^C NMR (175 MHz, DMSO-*d*_*6*_): δ 166.6 (C-2), 158.9 (C-5), 154.9 (C-3a), 154.4 (C-1′), 150.5 (C-9a), 146.0 (C-8), 138.8 (C-6), 131.5 (C-1′′), 130.4 (C-3′/5′), 125.7 (C-4′), 122.6 (C-5a), 120.1 (C-2′/6′), 118.0 (C-3′′), 113.1 (C-7), 46.0 (C-1′′); HRMS *m/z* (EI): [M]^•+^ calcd. for C_17_H_13_N_5_O_2_, 319.1069; found, 319.1099.

### 4-Benzyl-2-phenoxypyrido[3,2-*e*][1,2,4]triazolo[1,5,-*a*]pyrimidin-5(4*H*)-one (2c)

White amorphous powder (70%); mp: 195–197 °C; IR: 1673 (C=O) cm^−1^; ^1^H NMR (500 MHz, DMSO-*d*_*6*_): δ 8.85 (dd, J = 4.8, 1.7 Hz, 1H, H-8), 8.62 (dd, J = 7.8, 1.7 Hz, 1H, H-6), 7.62 (dd, J = 7.8, 4.8 Hz, 1H, H-7), 7.49 (t, J = 7.8 Hz, 2H, H-3′/5′), 7.46 (d, J = 7.8 Hz, 2H, H-2′′/6′′), 7.36 (d, J = 7.8 Hz, 2H, H-2′/6′), 7.34 (t, J = 7.8 Hz, 2H, H-3′′/5′′), 7.29 (m, H-4′, 2H, 4′′), 5.32 (s, 2H, H-7′′); ^13^C NMR (175 MHz, DMSO-*d*_*6*_): δ 166.5 (C-2), 159.3 (C-5), 155.0 (C-3a), 154.3 (C-1′), 150.7 (C-9a), 146.1 (C-8), 138.8 (C-6), 135.9 (C-1′′), 130.3 (C-3′/5′), 128.9 (C-3′′/5′′), 128.3 (C-2′′/6′′), 128.1 (C-4′′), 125.7 (C-4′), 122.6 (C-5a), 120.1 (C-2′/6′), 113.1 (C-7), 47.2 (C-7′′); HRMS *m/z* (EI): [M]^•+^ calcd. for C_21_H_15_N_5_O_2_, 369.1226; found 369.1276.

### 4-(2-Methylbenzyl)-2-phenoxypyrido[3,2-*e*][1,2,4]triazolo[1,5,-*a*]pyrimidin-5(4*H*)-one (2d)

White amorphous powder (75%); mp: 160–162 °C; IR: 1670 (C=O) cm^−1^; ^1^H NMR (700 MHz, DMSO-*d*_*6*_): δ 8.87 (dd, J = 4.8, 1.7 Hz, 1H, H-8), 8.63 (dd, J = 7.8, 1.7 Hz, 1H, H-6), 7.64 (dd, J = 7.8, 4.8 Hz, 1H, H-7), 7.48 (t, J = 7.9 Hz, 2H, H-3′/5′), 7.34 (d, J = 7.9 Hz, 2H, H-2′/6′), 7.29 (t, J = 7.9 Hz, 1H, H-4′), 7.22 (d, J = 7.4 Hz, 1H, H-6′′), 7.18 (m, 2H, H-3′′/5′′), 7.08 (t, J = 7.4 Hz, 1H, H-4′′), 5.28 (s, 2H, H-7′′), 2.42 (s, 3H, Ar-*Me*); ^13^C NMR (175 MHz, DMSO-*d*_*6*_): δ 166.5 (C-2), 159.4 (C-5), 155.0 (C-3a), 154.3 (C-1′), 150.8 (C-9a), 146.2 (C-8), 138.9 (C-6), 135.8 (C-1′′), 133.8 (C-2′′), 130.5 (C-3′′), 130.3 (C-3′/5′), 127.7 (C-4′′), 126.5 (C-5′′), 126.4 (C-6′′), 125.7 (C-4′), 122.6 (C-5a), 120.1 (C-2′/6′), 113.1 (C-7), 45.1 (C-7′′), 19.4 (Ar-*Me*); HRMS *m/z* (EI): [M]^•+^ calcd. for C_22_H_17_N_5_O_2_, 383.1382; found 383.1412.

### 4-(3-Methylbenzyl)-2-phenoxypyrido[3,2-*e*][1,2,4]triazolo[1,5,-*a*]pyrimidin-5(4*H*)-one (2e)

White amorphous powder (79%); mp 181–183 °C; IR: 1678 (C=O) cm^−1^; ^1^H NMR (700 MHz, DMSO-*d*_*6*_): δ 8.87 (d, J = 4.0 Hz, 1H, H-8), 8.64 (d, J = 7.6 Hz, 1H, H-6), 7.65 (dd, J = 7.8, 4.8 Hz, 1H, H-7), 7.51 (t, J = 7.8 Hz, 2H, H-3′/5′), 7.38 (d, J = 7.8 Hz, 2H, H-2′/6′), 7.32 (t, J = 7.8 Hz, 1H, H-4′), 7.28 (br s, 1H, H-2′′), 7.25 (d, J = 7.4 Hz, 1H, H-6′′), 7.23 (t, J = 7.4 Hz, 1H, H-5′′), 7.11 (d, J = 7.4 Hz, 1H, H-4′′), 5.29 (s, 2H, H-7′′), 2.29 (s, 3H, Ar-*Me*); ^13^C NMR (175 MHz, DMSO-*d*_*6*_): δ 166.5 (C-2), 159.3 (C-5), 155.0 (C-3a), 154.4 (C-1′), 150.7 (C-9a), 146.1 (C-8), 138.9 (C-6), 138.1 (C-1′′), 135.9 (C-3′′), 130.3 (C-3′/5′), 128.8 (C-2′′), 128.7 (C-5′′), 128.0 (C-4′′), 125.7 (C-4′), 125.4 (C-6′′), 122.6 (C-5a), 120.1 (C-2′/6′), 113.1 (C-7), 47.2 (C-7′′), 21.5 (Ar-*Me*); HRMS *m/z* (EI): [M]^•+^ calcd. for C_22_H_17_N_5_O_2_, 383.1382; found 383.1412.

### 4-(3-Methoxybenzyl)-2-phenoxypyrido[3,2-*e*][1,2,4]triazolo[1,5,-*a*]pyrimidin-5(4*H*)-one (2f)

White amorphous powder (83%); mp: 189–191 °C; IR: 1682 (C=O) cm^−1^; ^1^H NMR (500 MHz, DMSO-*d*_*6*_): δ 8.85 (dd, J = 4.7, 1.5 Hz, 1H, H-8), 8.63 (dd, J = 7.8, 1.5 Hz, 1H, H-6), 7.62 (dd, J = 7.8, 4.7 Hz, 1H, H-7), 7.48 (t, J = 7.8 Hz, 2H, H-3′/5′), 7.36 (d, J = 7.8 Hz, 2H, H-2′/6′), 7.29 (t, J = 7.4 Hz, 1H, H-4′), 7.25 (t, J = 8 Hz, 1H, H-5′′), 7.01 (m, 2H, H-2′′/6′′), 6.86 (dd, J = 8, 1.6 Hz, 1H, H-4′′), 5.28 (s, 2H, H-7′′), 3.72 (s, 3H, O-*Me*); ^13^C NMR (175 MHz, DMSO-*d*_*6*_): δ 166.5 (C-2), 159.8 (C-3′′), 159.3 (C-5), 154.9 (C-3a), 154.3 (C-1′), 150.7 (C-9a), 146.1 (C-8), 138.8 (C-6), 137.5 (C-1′′), 130.3 (C-3′/5′), 130.0 (C-5′′), 125.7 (C-4′), 122.6 (C-5a), 120.4 (C-6′′), 120.1 (C-2′/6′), 114.1 (C-4′′), 113.3 (C-7), 113.1 (C-2′′), 47.1 (C-7′′), 55.5 (O-*Me*); HRMS *m/z* (EI): [M]^•+^ calcd. for C_22_H_17_N_5_O_3_, 399.1331; found 399.1374.

### 4-((5-Oxo-2-phenoxypyrido[3,2-*e*][1,2,4]triazolo[1,5,-*a*]pyrimidin-4(5*H*)-yl)methyl)benzo-nitrile (2g)

White amorphous powder (73%); mp: 149–151 °C; IR: 1688 (C=O) cm^−1^; ^1^H NMR (700 MHz, DMSO-*d*_*6*_): δ 8.86 (dd, J = 4.9, 1 Hz, 1H, H-8), 8.62 (dd, J = 7.8, 1 Hz, 1H, H-6), 7.82 (d, J = 8.1 Hz, 2H, H-3′′/5′′), 7.66 (d, J = 8.1 Hz, 2H, H-2′′/6′′), 7.63 (dd, J = 7.8, 4.9 Hz, 1H, H-7), 7.48 (t, J = 7.8 Hz, 2H, H-3′/5′), 7.34 (d, J = 7.8 Hz, 2H, H-2′/6′), 7.29 (t, J = 7.8 Hz, 1H, H-4′), 5.39 (s, 2H, H-7′′); ^13^C NMR (175 MHz, DMSO-*d*_*6*_): δ 166.4 (C-2), 159.4 (C-5), 155.0 (C-3a), 154.3 (C-1′), 150.6 (C-9a), 146.2 (C-8), 141.6 (C-1′′), 138.8 (C-6), 133.2 (C-3′′/5′′), 130.3 (C-3′/5′), 128.9 (C-2′′/6′′), 125.7 (C-4′), 122.6 (C-5a), 120.1 (C-2′/6′), 119.2 (-C=N), 113.2 (C-7), 110.8 (C-4′′), 46.9 (C-7′′); HRMS *m/z* (EI): [M]^•+^ calcd. for C_22_H_14_N_6_O_2_, 394.1178; found 394.1211.

### 4-(4-Nitrobenzyl)-2-phenoxypyrido[3,2-*e*][1,2,4]triazolo[1,5,-*a*]pyrimidin-5(4*H*)-one (2h)

White amorphous powder (77%); mp: 150–152 °C; IR: 1672 (C=O) cm^−1^; ^1^H NMR (700 MHz, DMSO-*d*_*6*_): δ 8.87 (d, J = 4.9, 1 Hz, 1H, H-8), 8.63 (d, J = 7.8, 1 Hz, 1H, H-6), 8.19 (d, J = 8.5 Hz, 2H, H-3′′/5′′), 7.74 (d, J = 8.5 Hz, 2H, H-2′′/6′′), 7.64 (dd, J = 7.8, 4.9 Hz, 1H, H-7), 7.48 (t, J = 7.7 Hz, 2H, H-3′/5′), 7.34 (d, J = 8.2 Hz, 2H, H-2′/6′), 7.29 (t, J = 7.3 Hz, 1H, H-4′), 5.45 (s, 2H, H-7′′); ^13^C NMR (175 MHz, DMSO-*d*_*6*_): δ 166.4 (C-2), 159.4 (C-5), 155.0 (C-3a), 154.3 (C-1′), 150.6 (C-9a), 147.4 (C-4′′), 146.2 (C-8), 143.7 (C-1′′), 138.8 (C-6), 129.3 (C-2′′/6′′), 130.3 (C-3′/5′), 125.8 (C-4′), 123.9 (C-3′′/5′′), 122.6 (C-5a), 120.1 (C-2′/6′), 113.1 (C-7), 46.8 (C-7′′); HRMS *m/z* (EI): [M]^•+^ calcd. for C_21_H_14_N_6_O_4_, 414.1077; found 414.1110.

### 4-(4-Chlorobenzyl)-2-phenoxypyrido[3,2-*e*][1,2,4]triazolo[1,5,-*a*]pyrimidin-5(4*H*)-one (2i)

White amorphous powder (70%); mp: 144–146 °C; IR: 1677 (C=O) cm^−1^; ^1^H NMR (700 MHz, DMSO-*d*_*6*_): δ 8.85 (dd, J = 4.5, 1.1 Hz, 1H, H-8), 8.62 (d, J = 7.8 Hz, 1H, H-6), 7.62 (dd, J = 7.6, 4.8 Hz, 1H, H-7), 7.49 (m, H-3′/5′, 4H, 3′′/5′′), 7.39 (d, J = 8.3 Hz, 2H, H-2′′/6′′), 7.35 (d, J = 8.1 Hz, 2H, H-2′/6′), 7.30 (t, J = 7.4 Hz, 1H, H-4′), 5.30 (s, 2H, H-7′′); ^13^C NMR (175 MHz, DMSO-*d*_*6*_): δ 166.5 (C-2), 159.3 (C-5), 155.0 (C-3a), 154.3 (C-1′), 150.6 (C-9a), 146.1 (C-8), 138.8 (C-6), 135.0 (C-1′′), 132.7 (C-4′′), 130.3 (C-3′/5′, 2′′/6′′), 128.8 (C-3′′/5′′), 125.7 (C-4′), 122.6 (C-5a), 120.1 (C-2′/6′), 113.1 (C-7), 46.6 (C-7′′); HRMS m/z (EI): [M]^•+^ calcd. for C_21_H_14_ClN_5_O_2_, 403.0836; found 403.0872.

### 4-(2-Morpholinoethyl)-2-phenoxypyrido[3,2-*e*][1,2,4]triazolo[1,5,-*a*]pyrimidin-5(4*H*)-one (2j)

White amorphous powder (50%); mp: 169–171 °C; IR 1680 (C=O) cm^−1^; ^1^H NMR (700 MHz, DMSO-*d*_*6*_): δ 8.85 (d, J = 4.6 Hz, 1H, H-8), 8.63 (d, J = 7.8 Hz, 1H, H-6), 7.63 (dd, J = 7.6, 4.8 Hz, 1H, H-7), 7.49 (t, J = 7.7 Hz, 2H, H-3′/5′), 7.33 (d, J = 7.9 Hz, 2H, H-2′/6′), 7.29 (t, J = 7.4 Hz, 1H, H-4′), 4.24 (t, J = 7 Hz, 2H, CH_2_-8”), 3.51 (br s, 4H, H-3”/5”), 2.69 (t, J = 7 Hz, 2H, CH_2_-7”), 2.47 (br s, 4H, H-2”/6”); ^13^C NMR (175 MHz, DMSO-*d*_*6*_): δ 166.9 (C-2), 159.1 (C-5), 155.0 (C-3a), 154.3 (C-1′), 150.6 (C-9a), 146.7 (C-8), 138.8 (C-6), 130.3 (C-3′/5′), 125.7 (C-4′), 122.6 (C-5a), 120.1 (C-2′/6′), 113.7 (C-7), 66.6 (C-3”/5”), 55.1 (CH_2_-7”), 53.7 (C-2”/6”), 41.1 (CH_2_-8”); HRMS *m/z* (EI): [M]^•+^ calcd. for C_20_H_20_N_6_O_3_, 392.1597; found 392.1629.

### 2-Phenoxy-4-(2-(piperidin-1-yl)ethyl)pyrido[3,2-e][1,2,4]triazolo[1,5-a]pyrimidin-5(4*H*)-one (2k)

White amorphous powder (45%); mp: 200–202 °C; IR: 1679 (C=O) cm^−1^; ^1^H NMR (700 MHz, DMSO-*d*_*6*_): δ 8.82 (d, J = 4.3 Hz, 1H, H-8), 8.60 (d, J = 7.8 Hz, 1H, H-6), 7.60 (dd, J = 7.8, 4.3 Hz, 1H, H-7), 7.37 (t, J = 8.3 Hz, 2H, H-3′/5′), 7.33 (d, J = 7.7 Hz, 2H, H-2′/6′), 7.27 (t, J = 7.4 Hz, 1H, H-4′), 4.27 (t, J = 7 Hz, 2H, CH_2_-8”), 2.79 (t, J = 7 Hz, 2H, CH_2_-7”), 2.57 (m, 4H, H-2”/6”), 1.48 (m, 4H, H-3”/5”), 1.36 (m, 2H, H-4”); ^13^C NMR (175 MHz, DMSO-*d*_*6*_): δ 166.6 (C-2), 159.1 (C-5), 154.9 (C-3a), 154.6 (C-1′), 150.5 (C-9a), 146.7 (C-8), 138.7 (C-6), 130.3 (C-3′/5′), 125.7 (C-4′), 122.6 (C-5a), 120.1 (C-2′/6′), 113.7 (C-7), 54.9 (CH_2_-7”), 54.3 (C-2”/6”), 40.3 (CH_2_-8”), 25.5 (C3”/5”), 23.9 (C-4”); HRMS m/z (EI): [M]^•+^ calcd. For C_21_H_22_N_6_O_2_, 390.1804; found 390.1838.

### 2-(3-(5-Oxo-2-phenoxypyrido[3,2-*e*][1,2,4]triazolo[1,5-*a*]pyrimidin-4(5*H*)-yl)propyl)isoindo-line-1,3-dione (2l)

White amorphous powder (71%); mp: 155–157 °C; IR: 1670 (C=O) cm^−1^; ^1^H NMR (700 MHz, DMSO-*d*_*6*_): δ 8.82 (dd, J = 4.2, 1.1 Hz, 1H, H-8), 8.55 (dd, J = 7.8, 1.1 Hz, 1H, H-6), 7.82 (m, H-5”/6”, 4H, H4”/7”), 7.59 (dd, J = 7.8, 4.2 Hz, 1H, H-7), 7.47 (t, J = 7.8 Hz, 2H, H-3′/5′), 7.33 (d, J = 8.1 Hz, 2H, H-2′/6′), 7.28 (t, J = 7.5 Hz, 1H, H-4′), 4.17 (t, J = 7.1 Hz, 2H, CH_2_-8”), 3.71 (t, J = 7.1 Hz, 2H, CH_2_-10”), 2.16 (quint, J = 7.1 Hz, 2H, CH_2_-9”); ^13^C NMR (175 MHz, DMSO-*d*_*6*_): δ 168.4 (C-1”/3”), 166.6 (C-2), 159.2 (C-5), 154.8 (C-3a), 154.3 (C-1′), 150.5 (C-9a), 145.9 (C-8), 138.6 (C-6), 134.9 (C-3a”/7a”), 132.0 (C-5”/6”), 130.3 (C-3′/5′), 125.7 (C-4′), 123.5 (C-4”/7”), 122.6 (C-5a), 120.0 (C-2′/6′), 113.0 (C-7), 41.1 (CH_2_-8”), 35.7 (CH_2_-10”), 26.1 (CH_2_-9”); HRMS *m/z* (EI): [M]^•+^ calcd. for C_25_H_18_N_6_O_4_, 466.1390; found 466.1421.

### Procedure for the synthesis of 2-phenoxypyrido[3,2-e][1,2,4]triazolo[1,5-a]pyrimidine-5(4H)-thione (3)

A mixture of compound **1** (1 mmol) and phosphorus pentasulfide (1 mmol) was refluxed in absolute pyridine (10 mL) for 3 h. After cooling, the mixture was poured into ice/water, the resulting yellow solid was filtered, washed thoroughly with water and dried. Yield (90%); mp: 270–272 °C; IR: 1272 (C=S) cm^−1^; ^1^H NMR (500 MHz, DMSO-*d*_*6*_): δ 14.02 (br s, 1H, -NH), 8.92 (dd, J = 8.1, 1.7 Hz, 1H, H-6), 8.88 (dd, J = 4.6, 1.7 Hz, 1H, H-8), 7.63 (dd, J = 8.1, 4.6 Hz, 1H, H-7), 7.48 (td, J = 7.6, 2.3 Hz, 2H, H-3′/5′), 7.35 (d, J = 7.7 Hz, 2H, H-2′/6′), 7.29 (t, J = 7.4 Hz, 1H, H-4′); ^13^C NMR (175 MHz, DMSO-*d*_*6*_): δ 186.0 (C-5), 167.3 (C-2), 155.6 (C-3a), 154.5 (C-1′), 148.6 (C-9a), 143.2 (C-8), 141.6 (C-6), 130.3 (C-3′/5′), 125.7 (C-4′), 124.7 (C-5a), 123.1 (C-7), 120.2 (C-2′/6′); HRMS *m/z* (EI): [M]^•+^ calcd. for C_14_H_9_N_5_OS, 295.0528; found 295.0563.

### General procedure for the synthesis of 5-((3-methoxybenzyl)thio)-2-phenoxypyrido [3,2-e] [1,2,4] triazolo [1,5-a] pyrimidine (4)

At room temperature, potassium carbonate (1 mmol) was added portion wise over a period of 10 min to a stirred solution of **1** (0.5 mmol) in DMF (5 mL). After stirring for 10 min, 3-methoxybenzyl bromide (1.5 mmol) was added dropwise and the reaction mixture left to stir at room temperature for 20 h. The mixture was poured into ice/water, the resulting solid was filtered, washed with water and dried. Yield (68%); mp 149–151 °C; ^1^H NMR (500 MHz, DMSO-*d*_*6*_): *δ* 9.04 (d, J = 4.5 Hz, 1H, H-8), 8.68 (d, J = 7.9 Hz, 1H, H-6), 7.73 (dd, J = 7.9, 4.5 Hz, 1H, H-7), 7.49 (t, J = 7.7 Hz, 2H, H-3′/5′), 7.38 (d, J = 7.7 Hz, 2H, H-2′/6′), 7.30 (t, J = 7.4 Hz, 1H, H-4′), 7.25 (t, J = 8 Hz, 1H, H-5′′), 7.13 (br s, 1H, H-2′′), 7.08 (d, J = 7.4 Hz, 1H, H-6′′), 6.86 (d, J = 7.4 Hz, 1H, H-4′′), 4.66 (s, 2H, H-7′′), 3.74 (s, 3H, O-*Me*); ^13^C NMR (175 MHz, DMSO-*d*_*6*_): δ 168.5 (C-5), 167.8 (C-2), 159.8 (C-3′′), 156.4 (C-9a), 154.5 (C-3a), 153.4 (C-1′), 143.8 (C-8), 138.2 (C-6), 135.8 (C-1′′), 130.3 (C-3′/5′), 130.2 (C-5′′), 125.7 (C-4′), 122.7 (C-5a), 121.9 (C-7), 120.4 (C-2′/6′), 120.1 (C-6′′), 115.4 (C-4′′), 113.6 (C-2′′), 55.5 (O-*Me*), 34.1 (C-7′′); HRMS m/z (EI): [M]^•+^ calcd. For C_22_H_17_N_5_O_2_S, 415.1103; found 415.1136.

### Procedure for the synthesis of 5-chloro-2-phenoxypyrido[3,2-e][1,2,4]triazolo[1,5-a]pyrimidine (5)

Compound **1** (2 mmol) was refluxed with phosphorus oxychloride (2 mL) in benzene (14 mL) for 3 h. After evaporation of the solvent, the residue was treated with a saturated aqueous solution of potassium carbonate. The resulting solid was filtered, washed thoroughly with water and dried. The product was obtained as a white amorphous powder (70%); mp: 190–192 °C; ^1^H NMR (500 MHz, DMSO-*d*_*6*_): δ 9.16 (dd, J = 4.6, 1.6 Hz, 1H, H-8), 8.84 (dd, J = 8.3, 1.6 Hz, 1H, H-6), 7.89 (dd, J = 8.3, 4.6 Hz, 1H, H-7), 7.50 (t, J = 7.7 Hz, 2H, H-3′/5′), 7.39 (d, J = 7.7 Hz, 2H, H-2′/6′), 7.33 (t, J = 7.4 Hz, 1H, H-4′); ^13^C NMR (175 MHz, DMSO-*d*_*6*_): δ 168.8 (C-2), 157.4 (C-5), 156.7 (C-9a), 154.3 (C-3a), 152.5 (C-1′), 145.4 (C-8), 138.5 (C-6), 130.4 (C-3′/5′), 125.9 (C-4′), 123.8 (C-5a), 120.4 (C-2′/6′), 114.3 (C-7); HRMS *m/z* (EI): [M]^•+^ calcd. for C_14_H_8_ClN_5_O, 297.0417; found 297.0451.

### Procedure for the synthesis of compounds 6a–d

Compound **5** (1 mmol) was refluxed with the appropriate amine (5 mmol) in pyridine (10 mL) for 5–7 h. After cooling, the mixture was poured into ice/water, the resulting solid was filtered, washed with water and dried.

### 2-Phenoxy-*N*-phenylpyrido[3,2-e][1,2,4]triazolo[1,5-a]pyrimidin-5-amine (6a)

Brown amorphous powder (61%); mp: 251–253 °C; ^1^H NMR (500 MHz, DMSO-*d*_*6*_): δ 10.15 (br s, 1H, - NH), 9.14 (d, J = 7.8 Hz, 1H, H-6), 8.95 (d, J = 4.0 Hz, 1H, H-8), 7.86 (d, J = 7.8 Hz, 2H, H-2′′/6′′), 7.74 (dd, J = 8.1, 4.6 Hz, 1H, H-7), 7.46 (m, H-3′/5′, 4H, 3′′/5′′), 7.34 (d, J = 7.8 Hz, 2H, H-2′/6′), 7.27 (t, J = 7.4 Hz, 1H, H-4′), 7.22 (t, J = 7.4 Hz, 1H, H-4′′); ^13^C NMR (175 MHz, DMSO-*d*_*6*_): δ 168.4 (C-2), 155.2 (C-9a), 154.8 (C-5, 3a), 154.7 (C-1′), 145.6 (C-8), 138.9 (C-6), 135.4 (C-1′′), 130.2 (C-3′/5′), 129.1 (C-3′′/5′′), 125.4 (C-4′), 125.0 (C-4′′), 123.15 (C-2′′/6′′), 121.7 (C-7), 120.4 (C-2′/6′), 108.3 (C-5a); HRMS *m/z* (EI): [M]^•+^ calcd. for C_20_H_14_N_6_O, 354.1229; found 354.1264.

### 2-Phenoxy-N-(*p*-tolyl)pyrido[3,2-e][1,2,4]triazolo[1,5-a]pyrimidin-5-amine (6b)

Pale green amorphous powder (60%); mp: 162–164 °C; ^1^H NMR (700 MHz, DMSO-*d*_*6*_): δ 10.10 (br s, 1H, -NH), 9.12 (d, J = 8.2 Hz, 1H, H-6), 8.94 (d, J = 4.1 Hz, 1H, H-8), 7.73 (m, H-7, 3H, 2′′/6′′), 7.47 (t, J = 7.7 Hz, 2H, H-3′/5′), 7.34 (d, J = 8.3 Hz, 2H, H-2′/6′), 7.27 (t, J = 7.4 Hz, 1H, H-4′), 7.24 (d, J = 8.1 Hz, 2H, H-3′′/5′′), 2.33 (s, 3H, Ar-*Me*); ^13^C NMR (175 MHz, DMSO-*d*_*6*_): δ 168.4 (C-2), 155.2 (C-9a), 154.8 (C-5), 154.7 (C-3a), 154.6 (C-1′), 145.6 (C-8), 136.3 (C-6), 135.3 (C-1′′), 134.2 (C-4′′), 130.2 (C-3′/5′), 129.5 (C-3′′/5′′), 125.3 (C-4′), 123.2 (C-2′′/6′′), 121.6 (C-7), 120.4 (C-2′/6′), 108.3 (C-5a), 21.1 (Ar-*Me*); HRMS m/z (EI): [M]^•+^ calcd. for C_21_H_16_N_6_O, 368.1386; found 368.1416.

### *N*-(4-Methoxyphenyl)-2-phenoxypyrido[3,2-e][1,2,4]triazolo[1,5-a]pyrimidin-5-amine (6c)

Green amorphous powder (64%); mp: 174–176 °C; ^1^H NMR (700 MHz, DMSO-*d*_*6*_): δ 10.10 (br s, 1H, -NH), 9.09 (d, J = 8.2 Hz, 1H, H-6), 8.94 (d, J = 4.6 Hz, 1H, H-8), 7.72 (m, H-7, 3H, 2′′/6′′), 7.46 (t, J = 7.7 Hz, 2H, H-3′/5′), 7.33 (d, J = 7.8 Hz, 2H, H-2′/6′), 7.27 (t, J = 7.4 Hz, 1H, H-4′), 7.02 (d, J = 7.8 Hz, 2H, H-3′′/5′′), 3.79 (s, 3H, O-*Me*); ^13^C NMR (175 MHz, DMSO-*d*_*6*_): δ 168.4 (C-2), 156.8 (C-4′′), 155.3 (C-9a), 154.9 (C-5), 154.8 (C-3a), 154.7 (C-1′), 145.6 (C-8), 135.2 (C-6), 131.7 (C-1′′), 130.2 (C-3′/5′), 125.3 (C-4′), 125.1 (C-2′′/6′′), 121.6 (C-7), 120.4 (C-2′/6′), 114.3 (C-3′′/5′′), 108.3 (C-5a), 55.7 (O-*Me*); HRMS m/z (EI): [M]^•+^ calcd. for C_21_H_16_N_6_O_2_, 384.1335; found 384.1369.

### *N*-(4-Ethoxyphenyl)-2-phenoxypyrido[3,2-e][1,2,4]triazolo[1,5-a]pyrimidin-5-amine (6d)

Green amorphous powder (68%); mp: 154–156 °C; ^1^H NMR (700 MHz, DMSO-*d*_*6*_): δ 10.18 (br s, −1H, NH), 9.08 (d, J = 7.4 Hz, 1H, H-6), 8.93 (d, J = 4.1 Hz, 1H, H-8), 7.71 (m, H-7, 3H, 2′′/6′′), 7.46 (t, J = 7.7 Hz, 2H, H-3′/5′), 7.33 (d, J = 8.2 Hz, 2H, H-2′/6′), 7.27 (t, J = 7.3 Hz, 1H, H-4′), 6.99 (d, J = 7.8 Hz, 2H, H-3′′/5′′), 4.04 (q, J = 7 Hz, 2H, -*CH*_*2*_-CH_3_), 1.35 (t, J = 7 Hz, 3H, -CH_2_-*CH*_*3*_); ^13^C NMR (175 MHz, DMSO-*d*_*6*_): δ 168.4 (C-2), 156.1 (C-4′′), 155.2 (C-9a), 154.9 (C-5), 154.7 (C-3a/1′), 145.5 (C-8), 135.2 (C-6), 131.6 (C-1′′), 130.2 (C-3′/5′), 125.3 (C-4′), 125.0 (C-2′′/6′′), 121.6 (C-7), 120.4 (C-2′/6′), 114.7 (C-3′′/5′′), 108.2 (C-5a), 63.6 (-*CH*_*2*_-CH_3_), 15.2 (-CH_2_-*CH*_*3*_); HRMS *m/z* (EI): [M]^•+^ calcd. for C_22_H_18_N_6_O_2_, 398.1491; found 398.1524.

### Procedure for the synthesis of*N*-(2-phenoxypyrido[3,2-e][1,2,4]triazolo[1,5-a]pyrimidin-5-yl)hydroxylamine (6e)

Compound **5** (1 mmol) of was refluxed with hydroxylamine hydrochloride (3 mmol) in pyridine (10 mL) for 5 hrs. After cooling, the mixture was poured into ice/water, the resulting solid was collected, washed with water and dried. White amorphous powder (63%); mp: 249–251 °C; ^1^H NMR (500 MHz, DMSO-*d*_*6*_): δ 11.55 (br s, 1H, -OH), 10.80 (br s, 1H, -NH), 9.08 (dd, J = 4.4, 1.1 Hz, 1H, H-8), 8.27 (dd, J = 7.8, 1.1 Hz, 1H, H-6), 7.45 (t, J = 8.2 Hz, 2H, H-3′/5′), 7.35 (dd, J = 7.8, 4.4 Hz, 1H, H-7), 7.30 (d, J = 7.9 Hz, 2H, H-2′/6′), 7.25 (t, J = 7.4 Hz, 1H, H-4′); ^13^C NMR (175 MHz, DMSO-*d*_*6*_): δ 166.7 (C-2), 154.6 (C-9a/5), 151.1 (C-3a), 150.2 (C-1′), 143.9 (C-8), 139.8 (C-6), 130.2 (C-3′/5′), 125.4 (C-4′), 122.1 (C-7), 120.2 (C-2′/6′), 112.6 (C-5a); HRMS *m/z* (EI): [M]^•+^ calcd. for C_14_H_10_N_6_O_2_, 294.0865; found 294.0899.

### Procedure for the synthesis of 5-hydrazinyl-2-phenoxypyrido[3,2-e][1,2,4]triazolo[1,5-a]pyrimidine (6f)

Compound **5** (2 mmol) was refluxed with hydrazine hydrate (10 mmol) in ethanol (15 mL) for 3 hr. After cooling, the resulting solid was filtered, washed with water and dried. The product was obtained as a pale brown solid (74%); mp: 241–243 °C; ^1^H NMR (500 MHz, DMSO-*d*_*6*_): δ 8.82 (dd, J = 4.4, 1.1 Hz, 1H, H-8), 8.74 (d, J = 8.0 Hz, 1H, H-6), 7.58 (dd, J = 8.2, 4.4 Hz, 1H, H-7), 7.45 (t, J = 8.3 Hz, 2H, H-3′/5′), 7.33 (d, J = 7.8 Hz, 2H, H-2′/6′), 7.26 (t, J = 7.40 Hz, 1H, H-4′); ^13^C NMR (175 MHz, DMSO-*d*_*6*_): δ 168.2 (C-2), 155.7 (C-9a), 154.7 (C-3a/5), 154.3 (C-1′), 145.1 (C-8), 134.1 (C-6), 130.1 (C-3′/5′), 125.2 (C-4′), 121.4 (C-7), 120.2 (C-2′/6′), 106.7 (C-5a); HRMS *m/z* (EI): [M]^•+^ calcd. for C_14_H_11_N_7_O, 293.1025; found 293.1060.

### Procedure for the synthesis of compounds 6g and 6h

A mixture of **6f** (1 mmol) and 2-hydroxy-5-methoxy benzaldehyde (1 mmol) or 2-methoxy benzaldehyde (1 mmol) was refluxed in ethanol (10 mL) for 3–5 hrs. Then, the solvent was removed under reduced pressure and the resulting solid was filtered, washed with water and dried.

### (*E*)-4-Methoxy-2-((2-(2-phenoxypyrido[3,2-e][1,2,4]triazolo[1,5-a]pyrimidin-5-yl)hydrazono)methyl)phenol (6g)

Yellow amorphous powder (77%); mp: 220–222 °C; ^1^H NMR (700 MHz, DMSO-*d*_*6*_): δ 12.51 (br s, 1H, -NH), 10.61 (br s, 1H, -OH), 8.96 (br s, H-6, 2H, H-7′′), 8.74 (d, J = 4.5 Hz, 1H, H-8), 7.75 (dd, J = 7.9, 4.4, 1H, H-7), 7.48 (t, J = 8.3 Hz, 2H, H-3′/5′), 7.36 (d, J = 7.5 Hz, 2H, H-2′/6′), 7.28 (br s, 2H, H-4′, 6′′), 7.02 (d, J = 8.3 Hz, 1H, H-4′′), 6.93 (d, J = 8.3 Hz, 1H, H-3′′), 3.75 (s, 3H, O-*Me*); ^13^C NMR (175 MHz, DMSO-*d*_*6*_): δ 168.4 (C-2), 162.5 (C-5), 154.9 (C-9a), 154.8 (C-3a), 154.7 (C-1′), 153.2 (C-5′′), 152.7 (C-2′′), 152.6 (C-7′′), 145.9 (C-8), 135.9 (C-6), 130.2 (C-3′/5′), 125.4 (C-4′), 121.5 (C-7), 120.9 (C-4′′), 120.3 (C-2′/6′), 118.8 (C-1′′), 117.9 (C-6′′), 113.4 (C-3′′), 107.0 (C-5a), 55.9 (O-*Me*); HRMS *m/z* (EI): [M]^•+^ calcd. for C_22_H_17_N_7_O_3_, 427.1393; found 427.1426.

### (*E*)-5-(2-(2-Methoxybenzylidene)hydrazinyl)-2-phenoxypyrido[3,2-e][1,2,4]triazolo[1,5-a]pyrimidine (6h)

Yellow amorphous powder (72%); mp: 111–113 °C; ^1^H NMR (700 MHz, DMSO-*d*_*6*_): δ 12.04 (br s, 1H, -NH), 8.94 (br s, H-6, 2H, H-7′′), 8.98 (d, J = 4.5 Hz, 1H, H-8), 7.73 (dd, J = 7.9, 4.4, 1H, H-7), 7.53 (t, J = 7.5 Hz, 1H, H-4′′), 7.46 (m, 3H, H-3′/5′, 6′′), 7.36 (d, J = 7.9 Hz, 2H, H-2′/6′), 7.28 (t, J = 7.4 Hz, 1H, H-4′), 7.15 (d, J = 8.3 Hz, 1H, H-3′′), 7.04 (t, J = 7.5 Hz, 1H, H-5′′), 3.89 (s, 3H, O-*Me*); ^13^C NMR (175 MHz, DMSO-*d*_*6*_): δ 168.5 (C-2), 159.2 (C-5), 157.0 (C-9a, 2′′), 154.8 (C-3a), 154.7 (C-1′), 152.1 (C-7′′), 145.6 (C-8), 135.9 (C-6), 133.5 (C-4′′), 132.3 (C-6′′), 130.2 (C-3′/5′), 126.9 (C-5′′), 125.4 (C-4′), 121.6 (C-7), 121.2 (C-1′′), 120.3 (C-2′/6′), 112.5 (C-3′′), 106.9 (C-5a), 56.3 (O-*Me*); HRMS *m/z* (EI): [M]^•+^ calcd. for C_22_H_17_N_7_O_2_, 411.1444; found 411.1477.

### Procedure for the synthesis of*N*′-(2-phenoxypyrido[3,2-e][1,2,4]triazolo[1,5-a]pyrimidin-5-yl)isonicotinohydrazide (6i)

A mixture of **5** (0.5 mmol) and isoniazid (1.1 mmol) was refluxed in toluene (10 mL) for 4 h. After cooling, the resulting solid was collected by filtration, washed and dried. The product was obtained as a pale green amorphous powder (74%); mp: 163–165 °C; ^1^H NMR (500 MHz, DMSO-*d*_*6*_): δ 11.30 (br s, 1H, -NH), 9.00 (d, J = 4.3, 2H, H-3′′/5′′), 8.83 (d, J = 4.5 Hz, 1H, H-8), 8.55 (d, J = 7.8 Hz, 1H, H-6), 8.23 (d, J = 4.3 Hz, 2H, H-2′′/6′′), 7.89 (dd, J = 8, 4.5 Hz, 1H, H-7), 7.51 (t, J = 8.1 Hz, 2H, H-3′/5′), 7.39 (d, J = 8.1 Hz, 2H, H-2′/6′), 7.25 (t, J= 7.40 Hz, 1H, H-4′); ^13^C NMR (175 MHz, DMSO-*d*_*6*_): δ 168.7 (C-2), 166.9 (C-7′′), 155.5 (C-9a), 154.7 (C-3a) 154.6 (C-5), 154.3 (C-1′), 149.7 (C-3′′/5′′), 145.4 (C-8), 138.5 (C-6), 137.8 (C-1′′), 130.3 (C-3′/5′), 125.4 (C-4′), 122.4 (C-3′′/5′′), 122.0 (C-7), 120.2 (C-2′/6′), 113.7 (C-5a); HRMS *m/z* (EI): [M]^•+^ calcd. for C_20_H_14_N_8_O_2_, 398.1240; found 398.1273.

### α-Glucosidase inhibitory assay

#### Preparation of reagents

Phosphate-buffered saline pH 6.5 was prepared at 20 °C by dissolving sodium phosphate monobasic (1.4 g), sodium chloride (8 g), potassium chloride (0.2 g) and potassium dihydrogen phosphate (0.2 g) in deionized water (1 L). The solution was adjusted to pH 6.5 at 20 °C with 1 M sodium hydroxide (NaOH) or hydrochloric acid (HCl). A 20-mM solution of *para*-nitrophenyl-α-D-glucopyranoside (*p*-NPG substrate) was prepared by dissolving 6 mg of *p*-NPG in 10 mL of buffer solution. The α-glucosidase enzyme solution was prepared by dissolving α-glucosidase enzyme type 1 from Baker’s yeast (1 mg) in cold phosphate-buffered saline (1000 µL), pH 6.5; 50 µL of this solution was mixed into 12 mL of cold phosphate-buffered saline to give a concentration of 0.125 unit/mL. For the screening of samples, 200 µg/mL concentration was prepared in 100% dimethyl sulfoxide (DMSO). Then, two-fold serial dilutions with 5% DMSO were prepared in a 96-well microplate to give final concentrations of 100, 50, 25, 12.5, 6.25, 3.13 and 1.56 µg/ml. Acarbose was used as positive control and prepared at a concentration of 200 µg/ml in 100% DMSO. Then, two-fold serial dilutions with 5% DMSO were prepared in a 96-well microplate to give final concentrations of 100, 50, 25, 12.5, 6.25, 3.13 and 1.56 µg/ml.

#### Method

A volume of 10 µL of 5% DMSO was added to the 96 wells of the microplate. Then, 10 µL of the stock sample was added to the first well and mixed. Subsequently, 10 µL of the sample from the first well was taken and added to the second well and this procedure was repeated for the two-fold serial dilutions. Next, 20 µL of the α-glucosidase enzyme, 40 µL of phosphate buffered saline at pH 6.5 and 20 µL of deionized water were added to each well in the 96-well microtiter plate and mixed. The mixture was pre-incubated at 37 °C for 10 min. Then, 10 µL of 20 mM *p*-NPG solution was added to the mixture and the absorbance at 0 min. was measured at 405 nm. The reaction was incubated for 30 min. at 37 °C and the absorbance was then measured. For the negative control, the sample was replaced with 5% of DMSO and acarbose was used as positive control. Experiments were performed in triplicate. The inhibition activity of α-glucosidase was determined based on the described method by Ahmad et al., 2011^13^. The percent inhibition (%) of α-glucosidase inhibitory activity was calculated using the equation:

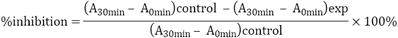

#### Statistical analysis

IC_50_ values are the mean values of three experiments represented as SEM (n=3). IC_50_ values are significantly different (P ≤ 0.05). Statistical analysis using one-way ANOVA was applied to the results. Nonlinear regression analysis using a four-parameter logistic curve was performed to calculate the IC_50_ values. The statistical software used were Microsoft Excel 2010 and Sigma Plot (version 14.0 Notebook, Systat Software Inc.).

#### Molecular docking

The Autodock software package was used to investigate the α-glucosidase inhibitory activity of the benzoquinazolines through molecular docking analysis^14^. The original docked protein were downloaded from the RCSB data bank web site (PDB file with code 3W37^15^). The detailed steps for the molecular docking studies were reported by Abuelizz and collaborators^16^.

## Acknowledgements

The authors extend their appreciation to the Deanship of Scientific Research at King Saud University for funding this work through research group No. RG-1439–011.

## Author contributions

R.A. and H.A.A. designed the work, the protocol of the study, synthesized-analyzed of compounds and wrote the manuscript. M.M. interpreted the NMR data and wrote the manuscript. R.A. and N. N. I. I. performed the biological studies and wrote their experimental data. A.E. performed and the molecular docking study. All the authors revised the whole manuscript.

## Corresponding author

Correspondence to Rashad Al-Salahi.

